# Machine Learning Models Reveal the Role of Ionization-Dependent Partitioning in Condensate Formation

**DOI:** 10.64898/2026.04.07.717090

**Authors:** Masoumeh Ozmaian, Seyyed Saeed Vaezzadeh

**Affiliations:** MatterMind Analytics LLC, Austin, 78731, TX; Lamar University, Beaumont, 77705, TX

## Abstract

Biomolecular condensates form through phase separation driven by multivalent interactions in eukaryotic cells, yet the factors that control small molecule partitioning remain incompletely understood. Building on previous evidence linking hydrophobicity and solubility to condensate affinity, we applied machine learning models to evaluate the role of ionization in this process. Using RDKit molecular descriptors, we trained regularized XGBoost regressors and classifiers across four representative condensates: cGAS-DNA, SUMO-SIM, SH3-PRM, and DHH1. Inclusion of logD, a pH dependent distribution coefficient that reflects effective lipophilicity, consistently improved predictive performance compared to models using only logP or logS. SHAP analysis identified logD as the dominant contributor to model predictions, suggesting that ionization coupled partitioning governs molecular localization within condensates. The addition of three-dimensional descriptors provided no further benefit, indicating that two dimensional physicochemical features and logD are sufficient to capture the main determinants of phase separation behavior. These findings establish logD as a mechanistic link connecting ionization, hydrophobicity, and small molecule partitioning in condensates, and offer a predictive framework for understanding small molecule behavior in these dynamic environments.

## Introduction

The cell interior is a highly crowded environment where biological macromolecules are distributed in a non-uniform manner. To manage this complexity, cells rely on the formation of biomolecular condensates, which arise through a variety of non-covalent interactions and help organize cellular components while supporting a wide range of biological processes. One well-studied mechanism behind this organization is complex coacervation, a type of liquid–liquid phase separation (LLPS) that drives the assembly of macromolecules into dynamic, membraneless compartments [1, 2]. This phenomenon underlies the formation of membraneless organelles (MLOs), which are essential for numerous biological functions. These highly dynamic condensates play diverse roles in cell physiology. For example, they act as reservoirs that regulate water availability and help maintain electrochemical balance within the cell [3-6]. In addition to electrostatic and hydrophobic forces, condensate formation is often driven by multivalent interactions, including π–π stacking between aromatic residues and cation–π interactions between arginine or lysine and aromatics, which provide an additional layer of selectivity and stability in phase separation. Besides their crucial role in cellular organization and function, biomolecular condensates have been implicated in diseases such as Alzheimer’s, Parkinson’s, type II diabetes, amyotrophic lateral sclerosis, and prion diseases[7, 8]. These disorders are linked to the pathological phase transitions of condensates, where dynamic, reversible liquid-like phases transform into irreversible, solid-like aggregates or fibrils. This solidification, driven by the imbalance of charged residues relative to aromatic and hydrophobic residues, disrupts cellular processes and contributes to neurodegeneration and disease progression, making it a key area of scientific research and therapeutic exploration [9-12].

Condensates can enrich small guest biomolecules either through partitioning, wherein molecules preferentially accumulate within the dense phase relative to the surrounding solution, or through interfacial adsorption, where they selectively bind to the condensate surface. Partitioning is not merely a passive accumulation of guest molecules within the dense phase; it can actively modulate condensate material properties such as viscosity, dynamics, and maturation. This selective enrichment may serve as a therapeutic mechanism by influencing condensate stability and modulating the kinetics of protein aggregation in both promoting and inhibitory directions[13]. Both the ability of a small molecule to associate with its target and the extent of its absorption are critical determinants of drug potency and represent key parameters in rational drug design. Unlike most drug targets that rely on specific binding sites, biomolecular condensates are dynamic assemblies of molecules that lack well-defined binding pockets, instead engaging guest molecules through nonspecific and multivalent interactions. This distinction underscores the need for large datasets to elucidate the mechanisms of molecular binding and small molecule uptake within condensates. Recent high-throughput efforts have been undertaken to investigate condensates, generating extensive measurements and data on the partitioning of molecules between the condensate interior and the surrounding solution, thereby providing insights into molecular enrichment and its potential functional consequences [14-22]. In the study by Thody et al [17], an extensive library of various small molecules and drug compounds were thoroughly examined on four different condensates and showed that solubility and hydrophobicity are the dominant indicators of partitioning. Ouyang et al [14] found that hydrophobic drugs were more effective in nonpolar residue-enriched condensates, and both binding affinity and hydrophobicity contributed significantly to enhancing inhibitor potency, highlighting the importance of condensate properties in drug design for condensates. Through extensive multi-scale molecular dynamics simulations, Emelianova et al. [15] also demonstrated that, in addition to physicochemical properties, small-molecule partitioning is also influenced by the affinity between the molecules and the condensate, suggesting the potential for selective partitioning.

Given the massive datasets typically generated in studies of small-molecule behavior and the substantial amount of information needed to elucidate therapeutic properties for drug design, data-driven models and machine learning algorithms offer powerful approaches for analyzing measurements and uncovering insights into phase separation and molecular partitioning [23-26]. Inspired by previous findings, we apply a range of Machine Learning (ML) algorithms to available experimental data [17] to systematically analyze the physicochemical properties and molecular descriptors of small molecules, with the goal of gaining deeper insight into the mechanisms that govern their partitioning behavior within biomolecular condensates. While our results confirm the importance of hydrophobicity and solubility reported in earlier studies, they also reveal that additional parameters, such as ionization state and pH dependent lipophilicity (captured by logD), play a critical role in modulating partitioning behavior across different condensate types. Our findings suggest that ionization coupled effects are central to small molecule localization within condensates and highlight logD as a robust and generalizable predictor of phase separation behavior, even in the absence of stereospecific interactions or three-dimensional structural features. Because condensates exhibit reduced dielectric environments relative to bulk solvent, the protonation state of small molecules directly modulates their desolvation penalty and transfer free energy, providing a mechanistic basis for the dominant role of logD.

## Results & Discussion

Figure 1 presents the SHAP-based feature importance profiles and corresponding beeswarm plots for models trained *without* inclusion of logD. Across all four condensates (cGAS-DNA, SUMO-SIM, SH3-PRM, and DHH1), the most influential descriptors are MolLogP (logP) and ESOL_LogS, followed by related hydrophobicity- and surface-area–based parameters.

**Figure 1.**
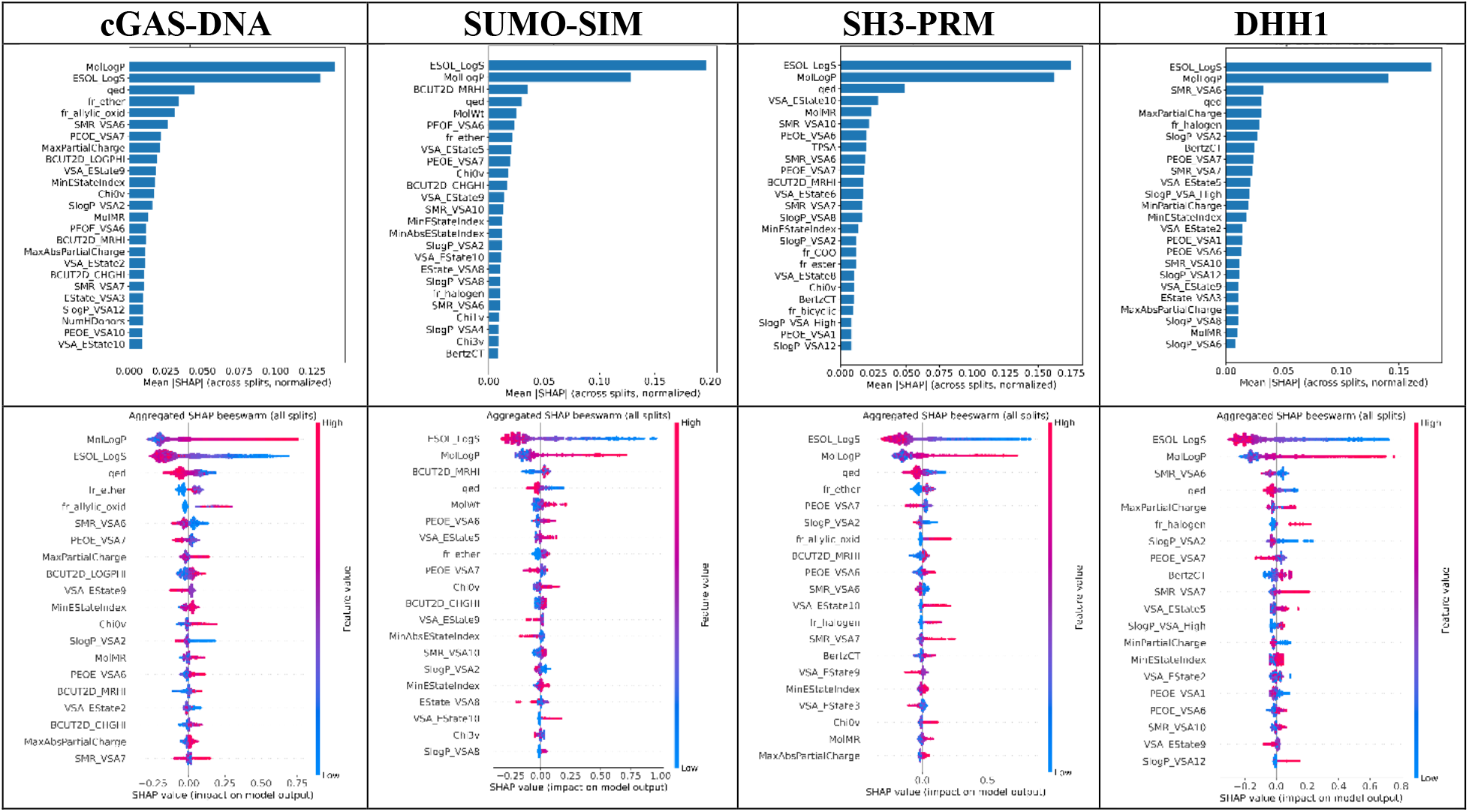
Feature importances (top) and beeswarm plots (bottom) of XGB models with all descriptors except for logD.

This pattern closely mirrors the findings reported by Thody et al. [27], confirming that lipophilicity and solubility are the primary physicochemical determinants of molecular partitioning into condensates. We therefore used this model configuration as a validation benchmark, demonstrating that our implementation faithfully reproduces the established feature hierarchy reported previously. The SHAP beeswarm plots further illustrate that molecules with higher logP values and lower predicted solubility (logS) are associated with increased model outputs, corresponding to stronger predicted partitioning into condensates.

Figure 2 summarizes the top 25 most influential features identified by SHAP analysis for the four condensate models (cGAS-DNA, SUMO-SIM, SH3-PRM, and DHH1) trained with inclusion of logD. In all condensates, introduction of logD markedly reshaped the feature importance landscape. LogD emerged as the single most influential descriptor, exceeding the contributions of MolLogP (logP) and ESOL_LogS (predicted aqueous solubility), which were the top-ranked predictors in the earlier “no logD” models. This finding indicates that pH-dependent partitioning, as captured by logD, provides additional explanatory power beyond static measures of lipophilicity (logP) or solubility (logS). Although logD dominates, logP and logS remain among the top-ranked features across all models, underscoring that baseline hydrophobicity and solvation energy remain central determinants of condensate partitioning. The simultaneous importance of these three descriptors (logD, logP, logS) highlights the interplay between static and dynamic solvation effects that govern biomolecular phase separation.

**Figure 2.**
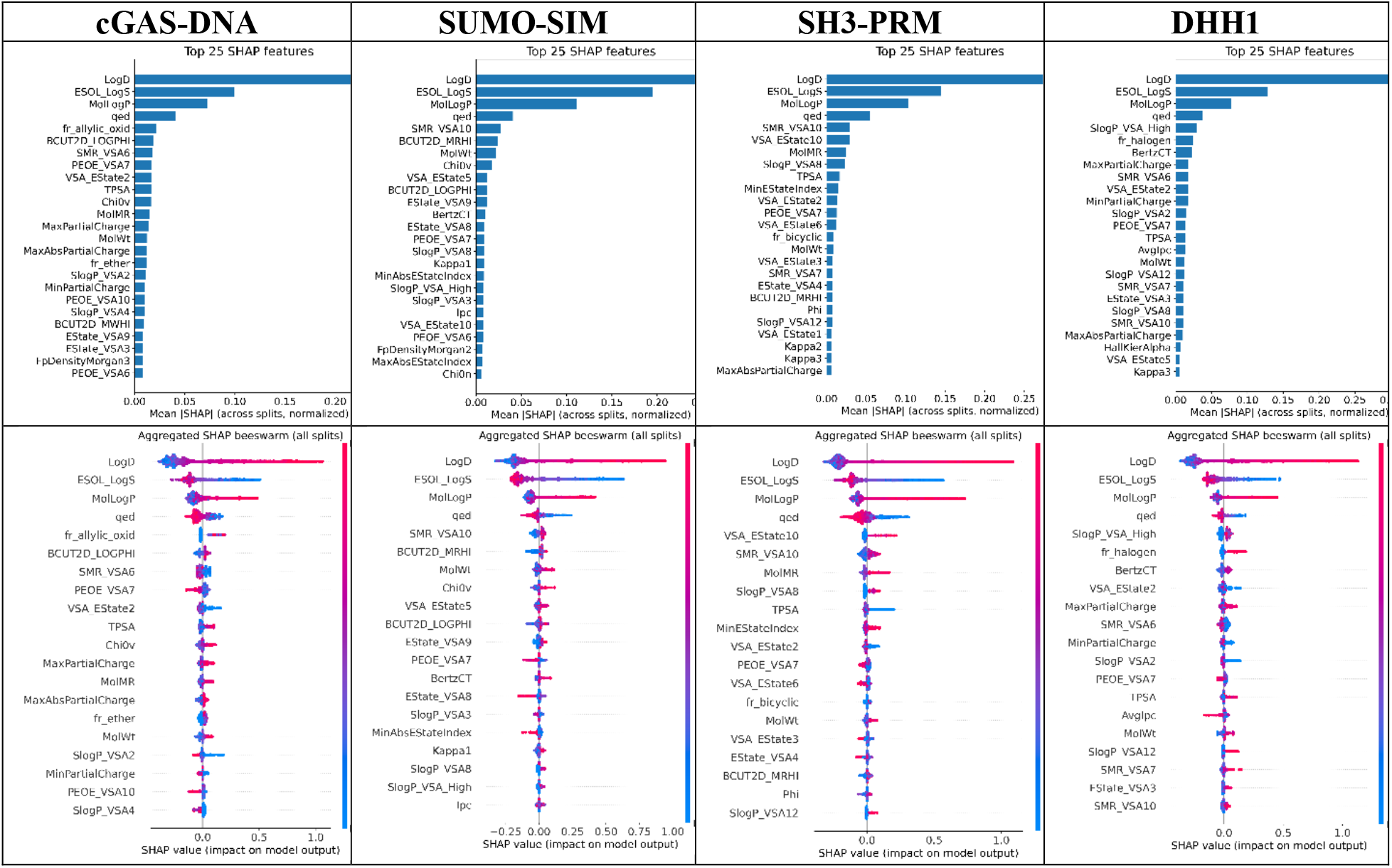
Feature importances (top) and beeswarm plots (bottom) of XGB models with all descriptors and including logD.

Across models, several other descriptors consistently appear among the top features: Quantitative Estimate of Drug-likeness (QED), a composite index that integrates multiple molecular properties such as molecular weight, lipophilicity, polar surface area, and hydrogen-bonding potential. Its prominence reflects the fact that condensate-partitioning molecules often share physicochemical profiles characteristic of drug-like compounds such as intermediate polarity, moderate size, and balanced solubility. SMR_VSA and SlogP_VSA descriptors, which quantify the molecular surface area associated with atomic fragments of particular refractivity or lipophilicity ranges. BCUT2D indices, which capture eigenvalue-based summaries of atomic charge and polarizability distributions, suggesting that electrostatic heterogeneity also contributes to condensate affinity.

Taken together, the SHAP profiles demonstrate that logD provides the dominant contribution to model accuracy while reinforcing the importance of molecular polarity, hydrophobic surface exposure, and charge distribution as secondary determinants of condensate partitioning.

Figure 3 illustrates the scatter plot of the mean partition coefficient across condensates (mean log PC) and logD. A clear positive correlation is observed, indicating that molecules with higher logD values, corresponding to greater lipophilic character at physiological pH, tend to partition more strongly into condensate phases. This trend supports the conclusion from our machine-learning analysis that ionization-dependent partitioning is a key determinant of condensate affinity. While other standard hydrophobicity descriptors such as log P capture only neutral-state lipophilicity, logD incorporates the effect of protonation and thus more accurately reflects the effective hydrophobicity under physiological conditions. The observed monotonic relationship between mean log PC and logD therefore provides direct experimental and computational evidence that dynamic, pH-dependent solvation behavior governs molecular partitioning into biomolecular condensates.

**Figure 3.**
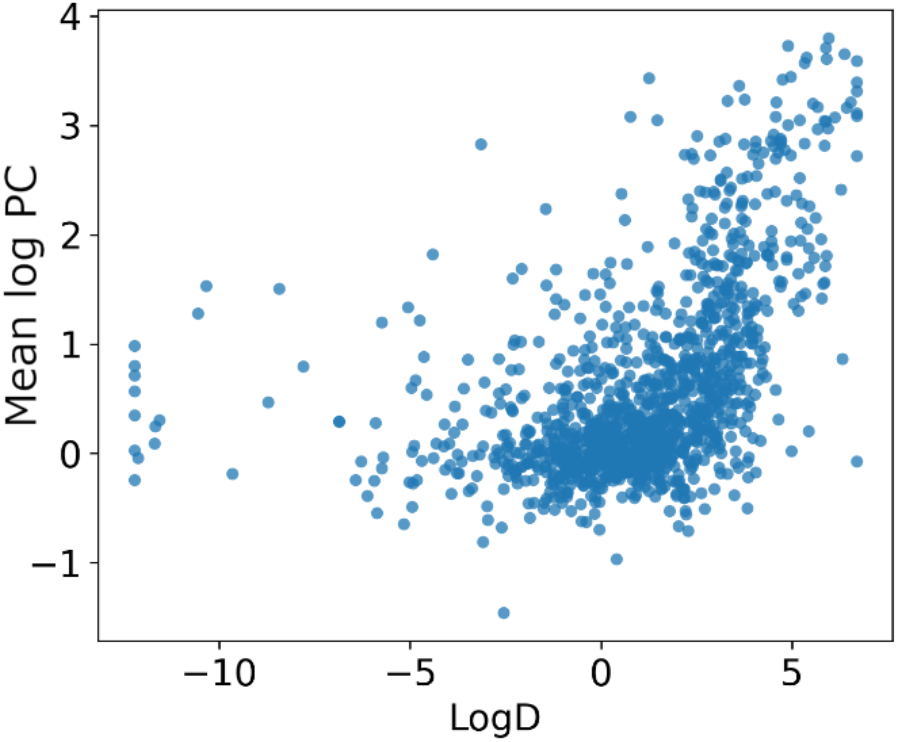
Dependence of mean log PC, averaged over all 4 condensates to logD

**Figure 4.**
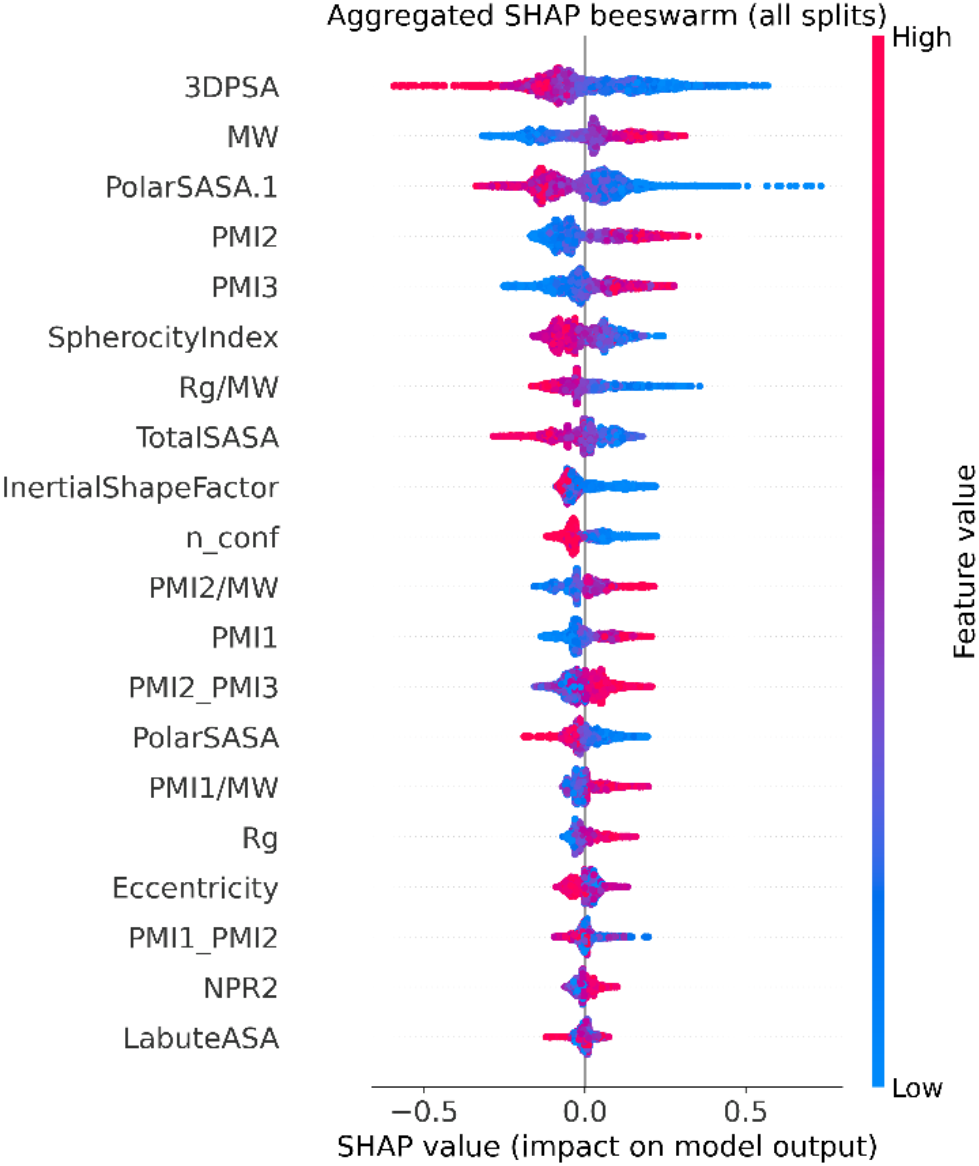
Beeswarm plot of SHAP summary for models including 3D geometric descriptors.

Table 1 summarizes the performance of XGBoost models trained to predict molecular partitioning across four condensates, with and without inclusion of logD as a feature. Among the four condensates analyzed, DHH1 and SH3-PRM showed the strongest dependence on inclusion of logD, exhibiting the largest gains in predictive performance (ΔR^2^ ≈ 0.06–0.07 and corresponding reductions in MAE and RMSE). cGAS-DNA and SUMO-SIM displayed smaller but consistent improvements, indicating that ionization-dependent partitioning captured by logD contributes most strongly to modeling condensate formation in DHH1 and SH3-PRM systems. Consistent with this trend, the improvement in predictive performance upon inclusion of logD was statistically significant across 20 train–test splits (paired t-test, p < 0.01).

**Table 1.** Performance of XGBoost models generated for different condensates as well as the corresponding average over all four condensates. The numbers show R2, MAE, and RMSE, respectively.

|  |  | cGAS-DNA | SUMO-SIM | SH3-PRM | DHH1 | Mean log PC |
| --- | --- | --- | --- | --- | --- | --- |
| no<br>logD | Train | 0.66/0.44/0.57 | 0.57/0.47/0.65 | 0.66/0.37/0.54 | 0.64/0.39/0.54 | 0.62/0.40/0.54 |
|  | Test | **0.52/0.52/0.67** | **0.40/0.55/0.77** | **0.55/0.44/0.64** | **0.50/0.45/0.62** | **0.45/0.46/0.63** |
| with<br>logD | Train | 0.67/0.42/0.56 | 0.57/0.46/0.65 | 0.68/0.36/0.52 | 0.68/0.36/0.50 | 0.64/0.38/0.53 |
|  | Test | **0.55/0.49/0.65** | **0.43/0.53/0.75** | **0.61/0.41/0.59** | **0.57/0.41/0.57** | **0.50/0.44/0.60** |

**Table 2.**
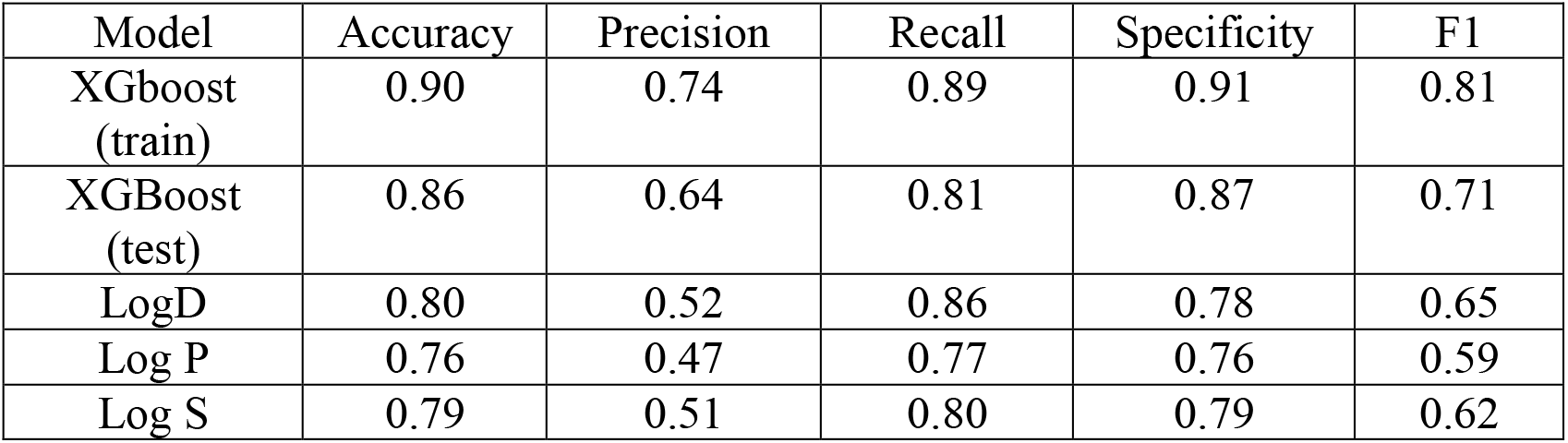
Classification performance comparison across identical data splits.

| Model | Accuracy | Precision | Recall | Specificity | F1 |
| --- | --- | --- | --- | --- | --- |
| XGboost (train) | 0.90 | 0.74 | 0.89 | 0.91 | 0.81 |
| XGBoost (test) | 0.86 | 0.64 | 0.81 | 0.87 | 0.71 |
| LogD | 0.80 | 0.52 | 0.86 | 0.78 | 0.65 |
| Log P | 0.76 | 0.47 | 0.77 | 0.76 | 0.59 |
| Log S | 0.79 | 0.51 | 0.80 | 0.79 | 0.62 |

### Assessment of 3D shape-based features

To evaluate whether three-dimensional molecular geometry contributes additional predictive power to condensate partitioning, we extended the feature set to include 3D shape and surface descriptors computed in RDKit. For each molecule, up to 200 conformers were generated using the ETKDG algorithm and energy-minimized with the MMFF94 force field. Descriptors were then computed for each conformer and averaged across all minimized conformations to obtain geometry-independent molecular statistics. The modeling protocol, including data splits, cross-validation, randomized hyperparameter optimization, and early stopping, was identical to that used for the 2D descriptor models to enable direct comparison.

The lack of improvement upon inclusion of 3D descriptors indicates that condensate partitioning is dominated by bulk physicochemical and solvation properties, which are already captured by 2D descriptors and logD.

Nonetheless, analysis of SHAP values (Figure X) reveals several 3D features that contribute moderately to prediction variance.

Descriptors such as 3DPSA (three-dimensional polar surface area) and PolarSASA quantify solvent-accessible polar surface, consistent with the role of molecular polarity and hydrogen-bonding potential in condensate formation.

The principal moments of inertia (PMI1–PMI3) and derived ratios (PMI2/PMI3, PMI1/PMI2, and SphericityIndex) capture molecular shape anisotropy, reflecting how elongated or compact geometries may influence solvation and multivalent interactions. The Radius of Gyration (Rg) and InertialShapeFactor similarly describe molecular compactness and mass distribution. We explicitly included Molecular Weight (MW) to decouple size-dependent effects from shape descriptors and confirm that the SHAP patterns are not dominated by trivial size correlations. Overall, these analyses suggest that while 3D geometric parameters provide mechanistic interpretability, highlighting the interplay between compactness, polarity, and surface exposure, they do not substantially improve predictive performance beyond 2D descriptors when trained under the same conditions.

### Binary classification of condensate partitioning

To complement the regression analysis, we trained a binary XGBoost classifier to distinguish molecules that partition into condensates from those that do not. The response was defined from the mean log PC across condensates using clear decision boundaries: compounds with 0.75≤log PC≤1.25 were excluded to avoid a gray zone around the threshold. We used stratified 80/20 train–test splits, repeated 10 times, preserving the empirical class ratio to minimize any effects of class imbalance. Also, the hyperparameter search, early stopping, and evaluation protocol mirrored the regression setup.

SHAP analysis of the classifier (Fig. 5) recapitulates the regression findings as expected: logD is the dominant predictor, followed by ESOL_LogS (log S), MolLogP (log P), and QED. We compared the full XGBoost classifier to single-feature baselines using only logD, only log S, or only log P. Receiver–operating characteristic (ROC) curves (Fig. 6) show that the full model achieves the highest AUC (0.901 ± 0.016), outperforming logD (0.872 ± 0.020), log S (0.845 ± 0.031), and log P (0.826 ± 0.037). Notably, logD alone is a strong baseline, approaching the multifeature model, consistent with its leading SHAP importance.

**Figure 5.**
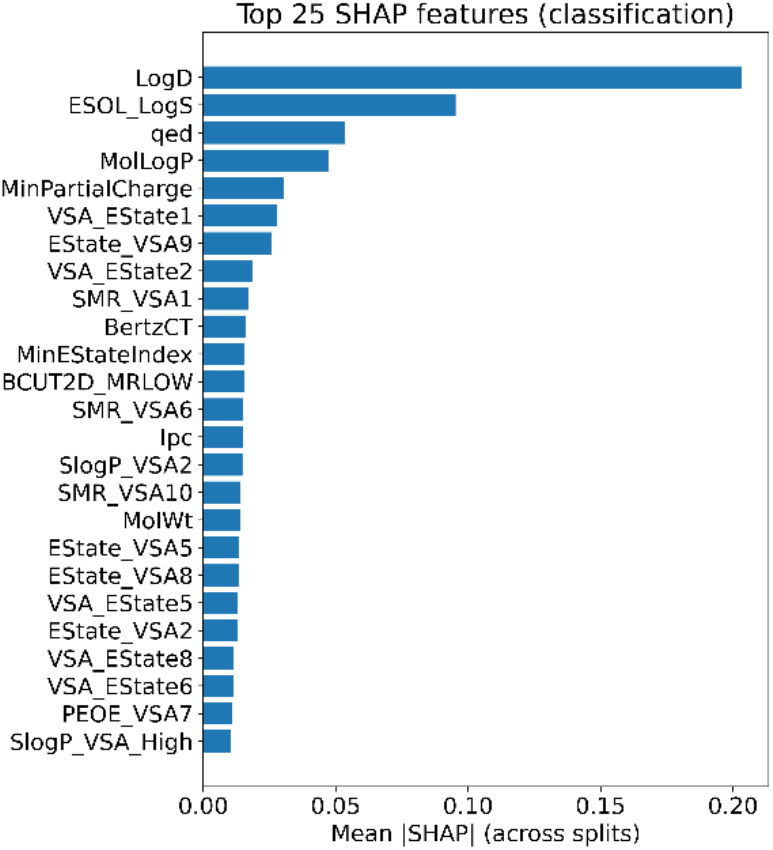
Top SHAP features for the binary classifier.

**Figure 6.**
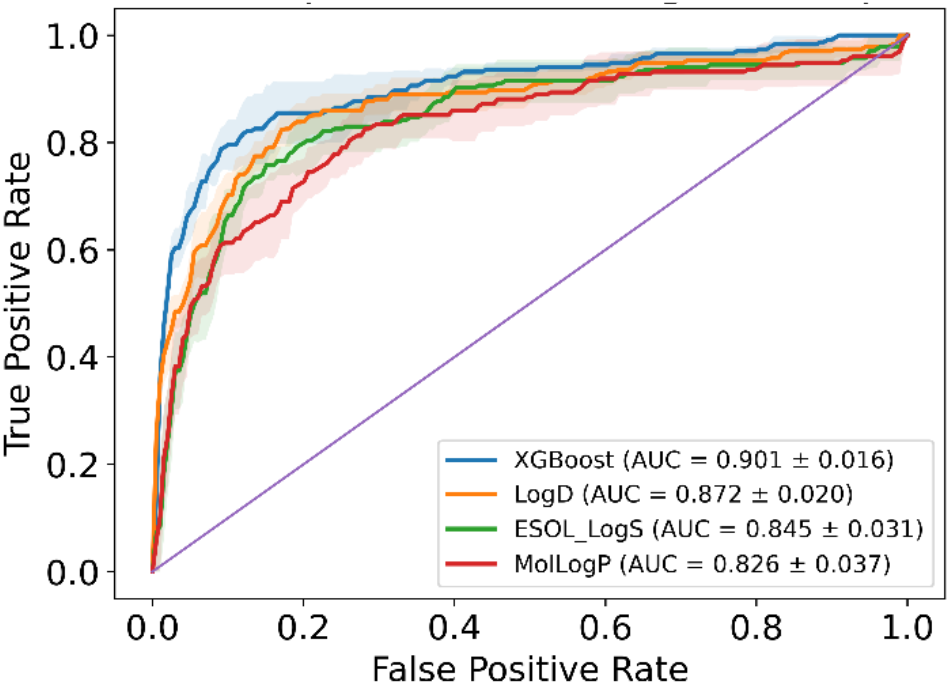
ROC curves comparing the full model to single-descriptor baselines. Mean ROC (shaded ±1 SD over 10 stratified 80/20 splits). XGBoost: AUC = 0.901 ± 0.016; logD: 0.872 ± 0.020; log S: 0.845 ± 0.031; log P: 0.826 ± 0.037. The multifeature model performs best overall, with logD alone a notably strong comparator.

To provide a quantitative comparison of classification performance, Table X reports the standard evaluation metrics for the XGBoost classifier and for single-descriptor baseline models using only logD, log P, or log S as inputs. To ensure a fair comparison between the multifeature classifier and single-descriptor baselines, all models were evaluated on the same stratified 80/20 train–test splits used for the XGBoost classifier. Each metric represents the mean value across ten randomized validation runs, maintaining consistent class balance and split composition across models.

The XGBoost classifier achieved the highest overall performance (test accuracy = 0.86, F_1_ = 0.71), with balanced recall = 0.81 and specificity = 0.87, indicating robust generalization. Among single-descriptor baselines, logD showed the strongest predictive power (accuracy = 0.80, F_1_ = 0.65), substantially outperforming log P and log S, which both exhibited weaker precision and specificity. These results reaffirm that ionization-dependent partitioning (logD) encapsulates most of the discriminative signal for condensate affinity, while the inclusion of broader physicochemical descriptors in the XGBoost model enhances precision and model stability.

## Conclusion

This study identifies ionization-dependent partitioning, quantified by logD, as a central determinant of small-molecule distribution within biomolecular condensates. Across all four systems examined, including logD as a feature markedly enhanced predictive accuracy relative to models relying solely on logP and logS, emphasizing that the effective lipophilicity of molecules under physiological pH conditions drives condensate affinity. SHAP analyses consistently ranked logD as the most influential descriptor, revealing a direct, monotonic relationship between logD and experimental partition coefficients.

Incorporating 3D structural descriptors provided complementary mechanistic insight into molecular geometry but did not further improve model accuracy, indicating that 2D physicochemical parameters and logD adequately represent the energetic determinants of condensate partitioning. Complementary classification models corroborated these findings, showing that logD alone captures most of the discriminative power for distinguishing partitioning versus non-partitioning compounds.

Collectively, these results establish a data-driven and interpretable framework for modeling small-molecule partitioning in condensates and highlight logD as a unifying descriptor connecting ionization, hydrophobicity, and phase-separation propensity—a principle that may guide rational design of chemical modulators targeting condensate-associated pathways. These findings suggest that tuning ionization equilibria (e.g., pKa) may provide a practical strategy for designing condensate-targeting small molecules.

## Method

### Molecular descriptors

For each compound, molecular descriptors were generated from the corresponding SMILES representation using RDKit. A comprehensive set of two-dimensional physicochemical descriptors was calculated, encompassing constitutional, topological, electronic, and surface area–related properties (208 descriptors). These descriptors capture intrinsic chemical features relevant to solubility, polarity, and molecular recognition.

To complement the RDKit baseline set with empirically motivated physicochemical parameters, we incorporated three additional descriptors: ESOL_LogS, an estimated aqueous solubility derived from the Delaney model (log mol/L), which encodes hydrophobic–hydrophilic balance and size effects, AromaticProportion, defined as the fraction of heavy atoms that are aromatic, capturing the degree of aromatic character and potential for π–π interactions, and SlogP_VSA_High, the cumulative van der Waals surface area associated with highly lipophilic atomic fragments (high-SlogP bins), reflecting hydrophobic surface exposure.

An explicit LogD term was included to account for ionization-dependent partitioning between hydrophobic and aqueous phases. This feature was calculated using ADMETlab3.0 [28]. LogD values were computed at pH 7.4 to match experimental partitioning conditions.

In addition to scalar physicochemical descriptors, 2D fingerprints were included in the model to capture substructural and topological molecular features. To this end Extended Connectivity Fingerprint of radius 2 (ECFP4, 2048 bit) is computed using RDKit.

### XGBoost models

All predictive modeling was performed using gradient-boosted decision trees. XGBoost was selected for its ability to model nonlinear relationships, handle correlated descriptors, and provide interpretable feature importances via both gain-based and SHAP analyses. Models were trained using the squared-error objective function, with the histogram-based tree method to accelerate training on large descriptor matrices. The primary optimization target was the coefficient of determination (R^2^) evaluated under internal cross-validation. To prevent overfitting and ensure robust generalization, model complexity was tightly regularized. The search space for hyperparameters was deliberately constrained to physically and statistically reasonable ranges for molecular property prediction:

- tree depth (max_depth): 1–3
- minimum child weight (min_child_weight): 20–35
- learning rate (η): 0.05–0.08
- L_1_ (α) and L_2_ (λ) regularization: α = 1–10, λ = 10–50
- subsampling of rows and columns (subsample, colsample_bytree): 0.5–0.7
- number of boosting rounds (n_estimators): 1500–3000
- structural penalty (gamma): 3.0–5.0
- histogram binning resolution (max_bin): 64–128

Hyperparameter optimization was conducted using randomized search cross-validation with five folds per search, exploring 450 randomized parameter sets. The best configuration was selected based on mean R^2^ across folds.

After hyperparameter selection, the model was retrained on each training subset with early stopping (patience of 25 rounds) using a 10% validation partition to determine the optimal number of boosting iterations. The final model was then refit on the complete training split using the best iteration count identified during early stopping.

Model performance was evaluated using 20 independent random 80/20 train–test splits, ensuring statistical robustness. For each split, we computed multiple regression quality metrics: root mean squared error (RMSE), mean absolute error (MAE), and R^2^ on both training and test sets.

Feature importance was extracted from SHapley Additive exPlanations (SHAP), which are based on cooperative game theory and quantify the contribution of each individual feature to a given prediction.

## REFERENCES

1. Abbas, M., et al., Peptide-based coacervate-core vesicles with semipermeable membranes. Advanced Materials, 2022. 34(34): p. 2202913.

2. Rai, S.K., et al., Heterotypic electrostatic interactions control complex phase separation of tau and prion into multiphasic condensates and co-aggregates. Proceedings of the National Academy of Sciences, 2023. 120(2): p. e2216338120.

3. Galvanetto, N., et al., Extreme dynamics in a biomolecular condensate. Nature, 2023. 619(7971): p. 876–883.

4. Watson, J.L., et al., Macromolecular condensation buffers intracellular water potential. Nature, 2023.

5. Uversky, V.N., Intrinsically disordered proteins in overcrowded milieu: Membrane-less organelles, phase separation, and intrinsic disorder. Current opinion in structural biology, 2017. 44: p. 18–30.

6. Dai, Y., et al., Biomolecular condensates regulate cellular electrochemical equilibria. Cell, 2024.

7. Nguyen, P.H., et al., Amyloid oligomers: A joint experimental/computational perspective on Alzheimer’s disease, Parkinson’s disease, type II diabetes, and amyotrophic lateral sclerosis. Chemical reviews, 2021. 121(4): p. 2545–2647.

8. Benson, M.D., et al., Amyloid nomenclature 2018: recommendations by the International Society of Amyloidosis (ISA) nomenclature committee. Amyloid, 2018. 25(4): p. 215–219.

9. Visser, B.S., W.P. Lipiński, and E. Spruijt, The role of biomolecular condensates in protein aggregation. Nature Reviews Chemistry, 2024: p. 1–15.

10. Erkamp, N.A., et al., Multiphase condensates from a kinetically arrested phase transition. bioRxiv, 2022: p. 2022.02. 09.479538.

11. Xia, Z., et al., Co-aggregation with Apolipoprotein E modulates the function of Amyloid-β in Alzheimer’s disease. Nature Communications, 2024. 15(1): p. 4695.

12. Silva, J.L., et al., Targeting biomolecular condensation and protein aggregation against cancer. Chemical reviews, 2023. 123(14): p. 9094–9138.

13. Lipiński, W.P., et al., Biomolecular condensates can both accelerate and suppress aggregation of α-synuclein. Science advances, 2022. 8(48): p. eabq6495.

14. Li, T., et al., Navigating condensate microenvironment to enhance small molecule drug targeting. 2024.

15. Emelianova, A., et al., Prediction of small-molecule partitioning into biomolecular condensates from simulation. JACS Au, 2025.

16. Mayfield, A., et al., Corelet™ platform: Precision high throughput screening for targeted drug discovery of biomolecular condensates. SLAS Discovery, 2025. 32: p. 100224.

17. Ambadi Thody, S., et al., Small-molecule properties define partitioning into biomolecular condensates. Nature Chemistry, 2024. 16(11): p. 1794–1802.

18. Kelley, F.M., et al., Controlled and orthogonal partitioning of large particles into biomolecular condensates. Nature communications, 2025. 16(1): p. 3521.

19. Klein, I.A., et al., Partitioning of cancer therapeutics in nuclear condensates. Science, 2020. 368(6497): p. 1386–1392.

20. Manzato, C., et al., Condensate screening identifies YM155 as β-catenin condensate inhibitor in colorectal cancer. bioRxiv, 2025: p. 2025.01. 13.632724.

21. Kilgore, H.R., et al., Distinct chemical environments in biomolecular condensates. Nature Chemical Biology, 2024. 20(3): p. 291–301.

22. Wang, C., et al., Nonspecific yet selective interactions contribute to small molecule condensate binding. Journal of chemical theory and computation, 2024. 20(22): p. 10247–10258.

23. van Mierlo, G., et al., Predicting protein condensate formation using machine learning. Cell reports, 2021. 34(5).

24. Saar, K.L., et al., Protein Condensate Atlas from predictive models of heteromolecular condensate composition. Nature Communications, 2024. 15(1): p. 5418.

25. Saar, K.L., et al., Learning the molecular grammar of protein condensates from sequence determinants and embeddings. Proceedings of the National Academy of Sciences, 2021. 118(15): p. e2019053118.

26. He, Y., et al., A High-Throughput, Flow Cytometry Approach to Measure Phase Behavior and Exchange in Biomolecular Condensates. bioRxiv, 2025: p. 2025.06.02.657082.

27. Ambadi Thody, S., et al., Small-molecule properties define partitioning into biomolecular condensates. Nature Chemistry, 2024: p. 1–9.

28. Fu, L., et al., ADMETlab 3.0: an updated comprehensive online ADMET prediction platform enhanced with broader coverage, improved performance, API functionality and decision support. Nucleic acids research, 2024. 52(W1): p. W422–W431.

